# Myoglianin triggers the pre-metamorphosis stage in hemimetabolan insects

**DOI:** 10.1101/368746

**Authors:** Orathai Kamsoi, Xavier Belles

## Abstract

Insect metamorphosis is triggered by a decrease in juvenile hormone (JH) in the final juvenile instar. What induces this decline is therefore a very relevant question. Working with the cockroach *Blattella germanica*, we found that Myoglianin (Myo), a ligand in the TGF-β signaling pathway, is highly expressed in the corpora allata (CA, the JH producing glands) and prothoracic glands (PG, which produce ecdysone) during the penultimate nymphal instar (N5). In the CA, high Myo levels during N5 repress the expression of *jhamt*, a JH biosynthesis gene. In the PG, decreasing JH levels trigger gland degeneration, mediated by the factors Kr-h1, FTZ-F1, E93 and IAP1. Also in the PG, a peak of *myo* expression in N5 stimulates the expression of ecdysone biosynthesis genes, such as *nvd*, thus enhancing the production of the metamorphic ecdysone pulse in N6. The *myo* expression peak in N5 also represses cell proliferation, which can contribute to enhance ecdysone production. The data indicate that Myo triggers the pre-metamorphic nymphal instar in *B. germanica*, and possibly in other hemimetabolan insects.

## INTRODUCTION

Insect metamorphosis has always been a fascinating phenomenon, but the master lines of their molecular regulatory mechanisms have only been elucidated recently. The process is regulated by two main hormones: ecdysone (where the best-known bioactive form is the derivative 20-hydroxyecdysone), which promotes the successive molts, and juvenile hormone (JH), which prevents the onset of metamorphosis in juvenile stages (Jindra et al., 2013; Nijhout, 1994). The molecular mechanisms underlying the action of these hormones are essentially based on the MEKRE93 pathway (Belles and Santos, 2014) that starts when JH binds to its receptor, Methoprene-tolerant (Met), which belongs to the bHLH-PAS family of transcription factors (Jindra et al., 2015b). Upon binding to Met, JH triggers dimerization of Met with another bHLH-PAS protein, Taiman (Tai). The resulting JH-Met+Tai complex induces transcription of the target gene *Krüppel-homolog 1* (*Kr-h1*), whose gene product represses the expression of *E93*, an ecdysone signaling-dependent gene whose gene product triggers metamorphosis (Belles and Santos, 2014; Jindra et al., 2015a).

In hemimetabolan metamorphosis, exemplified by the German cockroach, *Blattella germanica*, JH titers in the hemolymph rapidly decrease at the beginning of the final (sixth) nymphal instar (N6) (Treiblmayr et al., 2006). This is accompanied by a sudden drop of *Kr-h1* mRNA levels (Lozano and Belles, 2011), as a result of the decrease in JH and the action of microRNA miR-2, which scavenges *Kr-h1* transcripts (Belles, 2017; Lozano et al., 2015). Consequently, *E93* becomes de-repressed and its expression increases, which triggers the onset of metamorphosis (Belles and Santos, 2014). These observations and knowledge of the MEKRE93 pathway are robust, but lead to the following key question: what triggers the decline in JH at the beginning of N6? As we believe the answer lies in the penultimate nymphal stage (N5), we searched for candidate genes in a series of transcriptomes prepared and sequenced in our laboratory, which covered the entire ontogeny of *B. germanica* (Ylla et al., 2018). The analyses revealed that one of the most highly expressed genes in N5 is *myoglianin* (*myo*) (Ylla et al. 2018, figure 5E).

Myoglianin (Myo) was discovered in *Drosophila melanogaster* as a new member of the TGF-β signaling pathway and is closely related to the vertebrate muscle differentiation factor myostatin. Expression studies led to the detection of maternally-derived *myo* transcripts, *myo* expression in glial cells in mid-embryogenesis and, subsequently, in the developing somatic and visceral muscles and cardioblasts (Lo and Frasch, 1999). Myo is a ligand in the Activin branch of the TGF-β signaling pathway (Peterson and O’Connor, 2014). In the context of metamorphosis, Myo has been shown to play a crucial role in the remodeling of the mushroom bodies in the larva–pupa transition of *D. melanogaster*, an action mediated by increased expression of the *Ecdysone receptor B1* (*EcR-B1*) gene in neural tissues (Awasaki et al., 2011). Regarding hemimetabolan insects, in the corpora allata (CA, the JH producing glands) of the cricket *Gryllus bimaculatus*, Ishimaru et al. (2016) have shown that Myo inhibits the expression of *juvenile hormone acid methyl transferase* (*jhamt*), a gene coding for the last and crucial enzyme in the JH biosynthetic pathway. Depletion of this enzyme in the final nymphal instar triggers a decrease of JH production and commits the cricket to metamorphose (Ishimaru et al., 2016).

Using *B. germanica* as a model, we have found that Myo has functions beyond repressing the expression of *jhamt* in the CA during the transition from the penultimate to the final nymphal stage. We have also observed that Myo plays significant roles in the PG, which help induce this transition and ultimately lead to the onset of metamorphosis. Our data indicate that Myo is an essential factor in terms of triggering the pre-metamorphosis stage in *B. germanica* and possibly in other hemimetabolan insects, as it acts on the the CA and the PG, the glands that produce the most important hormones regulating metamorphosis.

## RESULTS

### Myoglianin structure and expression in *B. germanica*

We obtained a cDNA of 1,611 bp, comprising a complete open reading frame (ORF), by combining BLAST search in *B. germanica* genome, transcriptomic data and PCR. Its conceptual translation gave a 537 amino acid protein (Fig. S1) with significant similarity to the Myo sequence of other species, such as the fly *D. melanogaster* and the cricket *G. bimaculatus*, as described by Lo and Frasch (1999) and Ishimaru et al. (2016), respectively (Fig. S2). As in other Myo orthologs, the full-length *B. germanica* Myo sequence contained the canonical RXXR processing site and seven cysteines in the carboxy-terminal portion of the ORF, which would correspond to the mature processed protein (Figs. S1 and S2).

We were first interested in assessing whether *myo* was differentially expressed in different tissues, and we measured its expression in brain, corpora cardiaca–corpora allata (CC–CA) complex, prothoracic gland (PG), ovary, fat body, epidermis, and muscle tissues in 3-day-old fifth instar female nymphs (N5D3) of *B. germanica*. Results showed that PG showed the highest mRNA levels. Lower expression was measured in brain, CC–CA and muscle tissues, and the lowest expression was observed in fat body, ovary and epidermal tissues (Fig. 1A).

**Fig. 1.**
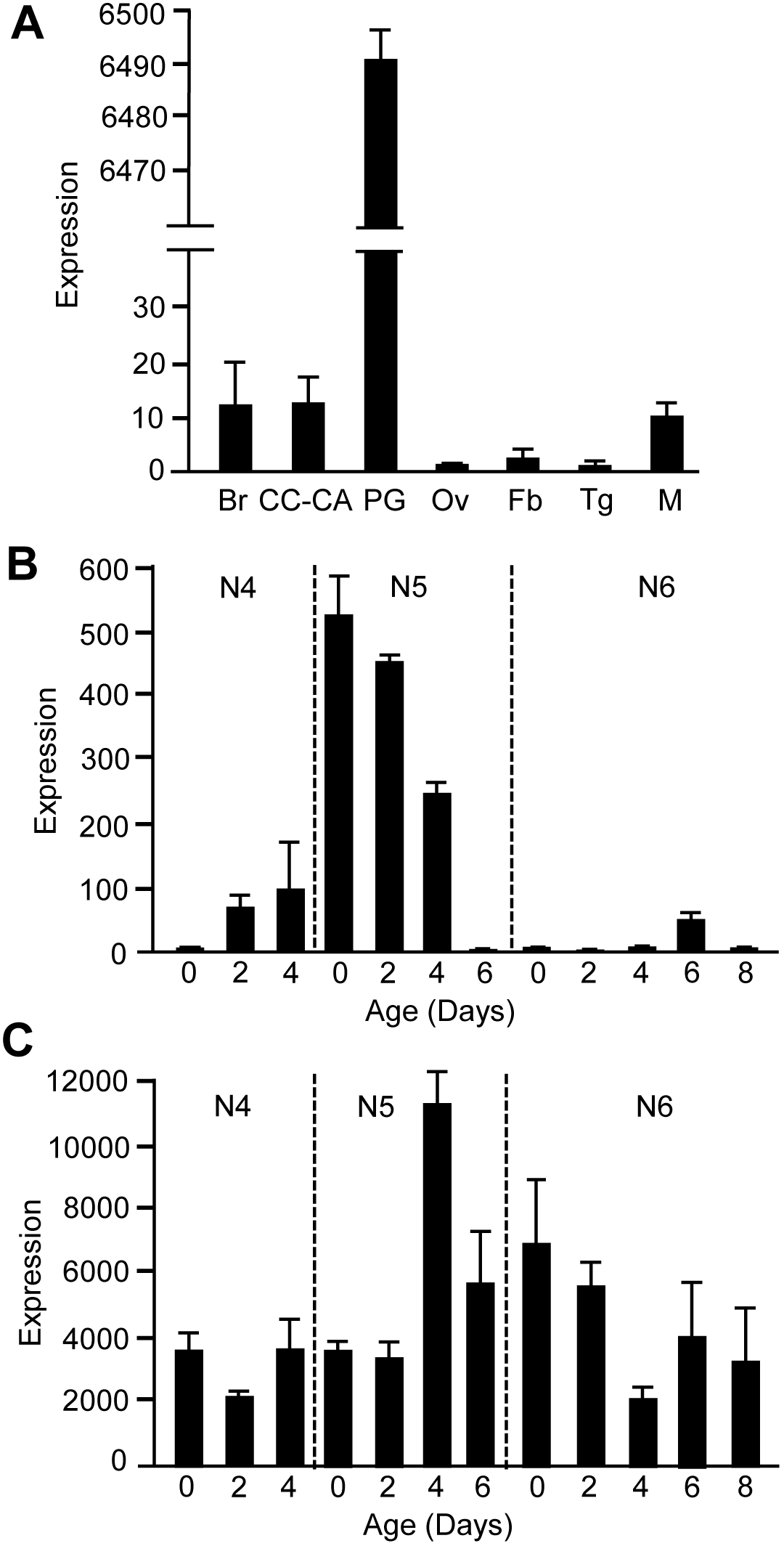
Expression of *myoglianin* in *Blattella germanica*. (A) mRNA levels in brain (Br), Corpora cardiaca-corpora allata complex (CC–CA), prothoracic gland (PG), ovary (Ov), fat body (Fb), epidermis (tergites two to seven: Tg), and muscle (extensor muscle from the six femora pairs: M); measurements were carried out on 3-day-old fifth instar female nymphs. (B) mRNA levels in CC–CA of fourth, fifth and sixth instars female nymphs. (C) mRNA levels in PG of fourth, fifth and sixth instars female nymphs. Results indicated as copies of *myoglianin* mRNA per 1000 copies of BgActin-5c mRNA and are expressed as the mean ± SEM (n=3-5).

As we were interested in the main hormones regulating metamorphosis, JH and ecdysone, we focused on the glands producing these hormones, the CC–CA complex and PG, respectively. Therefore, we determined the *myo* expression pattern in these two glands during the fourth, fifth, and sixth (final) instar female nymphs (N4, N5 and N6). In the CC–CA, the highest expression was observed in N5, in accordance with the pattern we found in our previous transcriptome analyses (Ylla et al., 2018). Within N5, maximal expression occurs at the beginning of the instar (N5D0) (ca. 500 mRNA copies per 1,000 copies of Actin mRNA) and progressively decreases until it practically vanishes at the end of the instar (N5D6). During N6, *myo* mRNA levels in the CC–CA are relatively low (between 10 and 50 mRNA copies per 1,000 copies of Actin mRNA) (Fig. 1B). In the PG, the highest *myo* mRNA levels were also observed in N5, but expression peaked in the mid-late instar (N5D4, ca. 11,000 mRNA copies per 1,000 copies of Actin mRNA). The levels then progressively decreased throughout the rest of the instar and N6, although they remained relatively high in N6 (around 4,000 mRNA copies per 1,000 copies of Actin mRNA) (Fig. 1C).

### Myoglianin depletion prevents metamorphosis

We used systemic RNAi to study the effects of Myo depletion on metamorphosis. We injected a single dose of 3 μg of a dsRNA targeting *myo* mRNA (dsMyo) into the abdomen of N5D0 cockroaches. Controls were treated equivalently with the same dose of an alien dsRNA (dsMock). In the CC–CA, *myo* mRNA levels were significantly lower in dsMyo-treated cockroaches (Fig. 2A). When monitoring the experimental cockroaches up to the imaginal molt, we observed that all N5 controls (n=44) molted to normal N6 after six days, on average, and then to normal adults eight days later (Fig. 2B). The N5 dsMyo-treated cockroaches (n=46) molted to N6 after eight days, on average; these N6 had an apparently normal morphology but were smaller than the N6 controls (Figs. 2C and D). Of the 46 N6 dsMyo-treated cockroaches, 6 died and 40 molted to a supernumerary nymph (N7). Of the 40 N7 nymphs, 6 molted to the adult stage and 34 molted to a further supernumerary nymph (N8), which then molted to adults. A total of 28 of these adults presented a practically normal morphology (albeit slightly larger than control adults emerging from N6), while the wings and tegmina were somewhat separated in the other 6 (Fig. 2D). The supernumerary nymphal instars N7 and N8 lasted for around eight days (Fig. 2B). We then examined the expression of genes involved in JH synthesis and signaling in the CC–CA at the beginning of N6. Results showed that the expression of *jhamt* and *Kr-h1* was significantly upregulated, whereas that of *E93* was downregulated (Fig. 2E).

**Fig. 2.**
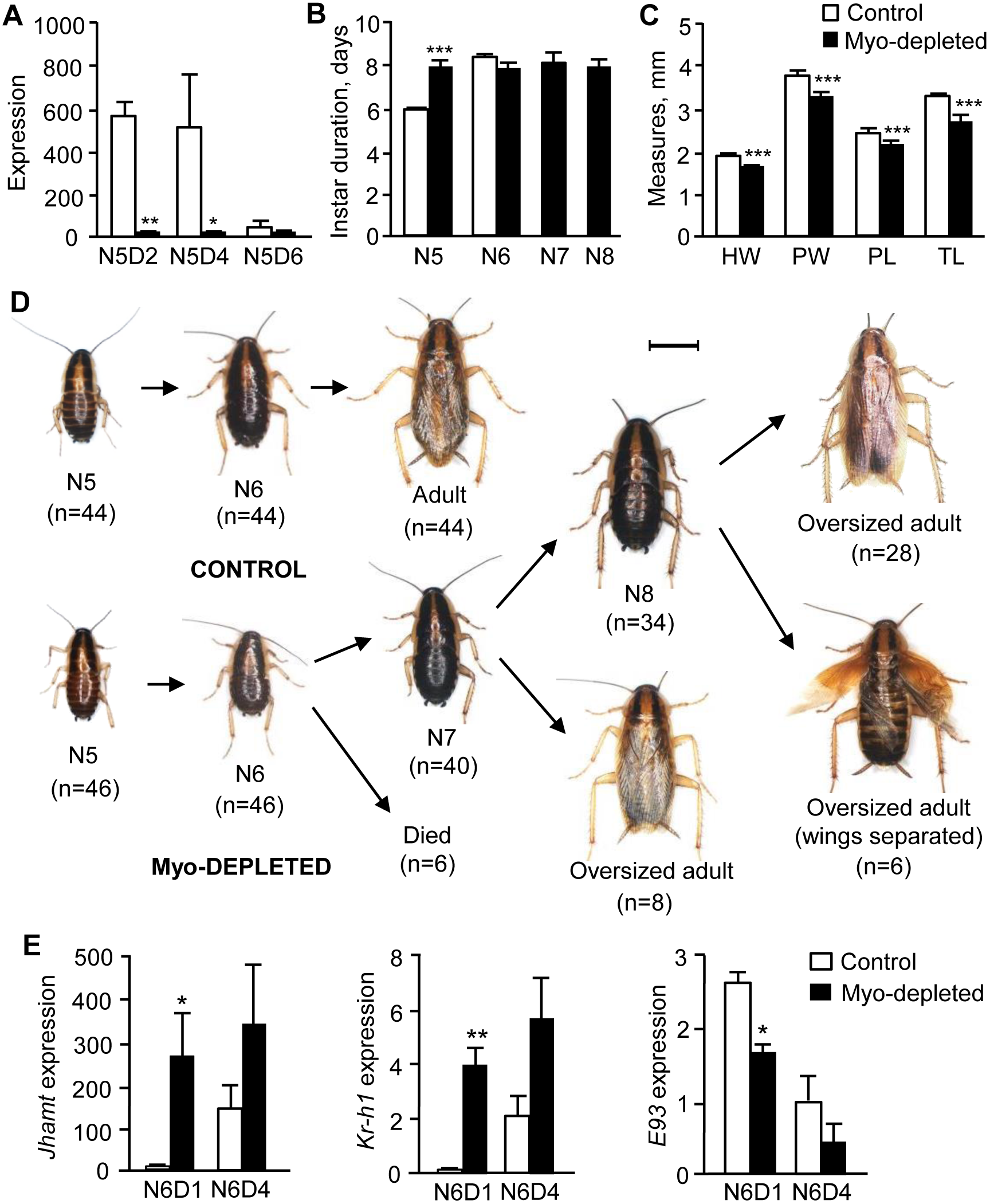
Effects of *myoglianin* mRNA depletion in the metamorphosis of *Blattella germanica*. (A) mRNA levels of *myo* in CC–CA of Myo-depleted cockroaches and in controls, measured on 2-, 4- and 6-day-old fifth instar female nymphs (N5D2, N5D4 and N5D6). (B) Duration (days) of N5 and N6 (and the supernumerary nymphal instars N7 and N8) in Myo-depleted cockroaches and in controls. (C) Measures of morphological parameters in N6 of Myo-depleted cockroaches and in controls; HW: head width, PW: pronotum width, PL: pronotum length, TL: metatibia length. (D) Dorsal view of control and Myo-depleted cockroaches through successive molts; freshly emerged fifth instar female nymphs (N5D0) were treated with dsMyo or with dsMock (control), and the morphology after the subsequent molts was examined; the number of specimens at the beginning of the experiments and after every molt is indicated; scale bar: 4 mm. (E) mRNA levels of *jhamt*, *Kr-h1* and *E93* in CC–CA from dsMyo-treated cockroaches and controls on N6D1 and N6D4. In A and E, results are indicated as copies of the given transcript per 1000 copies of BgActin-5c mRNA. In all bar diagrams, results are expressed as the mean ± SEM (n=3-5) and asterisks indicate statistically significant differences with respect to controls (*p<0.05, **p<0.01, ***p<0.001) according to the student’s *t*-test.

In the cricket *G. bimaculatus*, Ishimaru et al. (2016) reported that Myo represses the expression of *jhamt* in the CA. Thus, depletion of Myo in the final nymphal instar upregulated *jhamt* expression, hence JH production increased, and the crickets molted to supernumerary nymphs instead of adults (Ishimaru et al., 2016). The results obtained in *B. germanica* are similar to those observed in *G. bimaculatus* in terms of metamorphosis phenotypes and the up-regulation of *jhamt* expression after Myo depletion. This is not surprising as both species are hemimetabolan and closely related phylogenetically, that is, within the “orthopteroid” group of the superorder Polyneoptera (Misof et al., 2014). Our results, however, unveil the molecular mechanisms preventing metamorphosis, which are based on the MEKRE93 pathway (Belles and Santos, 2014). In the last nymphal instar, the *jhamt* up-regulation induced by Myo depletion led to an increase in *Kr-h1* expression, a JH-dependent factor that is the main transducer of the antimetamorphic action of JH (Lozano and Belles, 2011). Moreover, Kr-h1 represses *E93* (Belles and Santos, 2014), which is the master factor triggering metamorphosis (Belles and Santos, 2014; Ureña et al., 2014). In our Myo-depleted cockroaches, the up-regulation of *Kr-h1* expression explains the down-regulation of *E93* (Fig. 2E), whereas the observed reduction in *E93* expression explains why the cockroaches did not molt to the adult stage.

These observations are consistent with our previous work wherein we reported that the TGF-β/Activin signaling pathway contributes to the activation of *E93* expression and repression of *Kr-h1* expression during the metamorphosis of *B. germanica* (Santos et al., 2016). In the context of holometabolan metamorphosis, studies in *D. melanogaster* have shown that the TGF-β/Activin branch is involved in wing disc patterning (Peterson and O’Connor, 2013) and neuronal remodeling (Zheng et al., 2003). Moreover, Gibbens et al. (2011) have shown that when the TGF-β/Activin signaling pathway was blocked in the PG, then the last instar larvae of *D. melanogaster* were developmentally arrested, because they were unable to produce the large pulse of ecdysone needed for metamorphosis. It has been demonstrated that Myo signaling through the TGF-β/Activin pathway is crucial for mushroom body remodeling in the transition from the last larval stage to the pupa, as it up-regulates neuronal *EcR-B1* expression (Awasaki et al., 2011). The same study revealed that *myo*-defective mutants can molt until the last larval instar and prepupate, but they become arrested before head inversion (Awasaki et al., 2011), which is consistent with the results reported by Gibbens et al. (2011).

### Myoglianin depletion causes cell hyperproliferation in the prothoracic gland

The PG of *B. germanica*, like in other cockroaches, has an X-shaped morphology and its size varies during successive instars. Size variation roughly corresponds to the dynamics of ecdysone production as the most active glands have bigger secretory cells that the inactive glands (Romañá et al., 1995). In the present work, we also studied the dynamics of cell proliferation in the PG during N5 and N6 using ethynyl deoxyuridine (EdU) labeling. In N5, which lasts for 6 days, we observed intense cell division between days 0 and 3 (the first 50% of the instar). In N6, which lasts for 8 days, intense cell division only occurred over days 0 and 1 (the first 12.5% of the instar) (Fig. 3). During these instars, the ecdysone pulse in N5 peaks sharply at around day 4, whereas in N6 ecdysone production follows a longer, shallower peak over days 5, 6 and 7 (Cruz et al., 2003). The non-overlapping patterns of cell division and the ecdysone peak indicate that PG cells alternate between periods of cell proliferation and ecdysone synthesis, as occurs in other secretory glands, such as the CA (Chiang et al., 1995). Notably, the form of the ecdysone pulse varies along the different molts; wider pulses appear to be characteristic of metamorphic molts (Sakurai, 2005). We believe the wide ecdysone pulse in N6, which triggers the metamorphic molt, would require an earlier and radical arrest of cell division, similar to the one observed in our Edu labeling observations (Fig. 3).

**Fig. 3.**
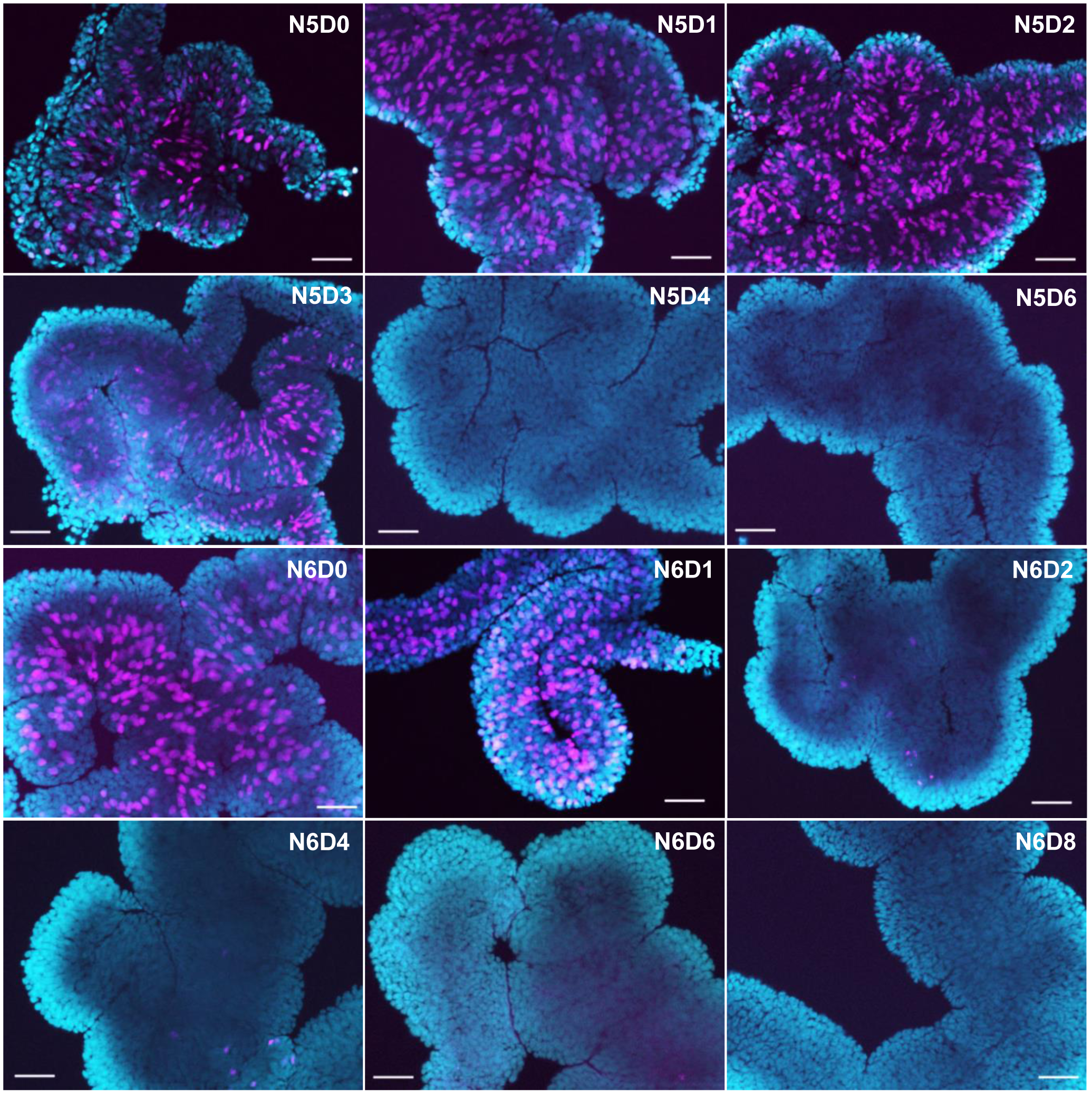
Cell proliferation in the prothoracic gland of *Blattella germanica*. *germanica*. (A) Double labeling EdU (discrete pink spots) and DAPI (background blue color) of PG tissue of female nymphs along the fifth and the sixth instar (from N5D0 to N5D6 and from N6D0 to N6D8); scale bar: 50 μm.

We used the same *myo* targeting RNAi experiment described above (3 μg of dsMyo injected in N5D0) to examine the effects in the PG. We found that *myo* mRNA levels in the PG were significantly lower in dsMyo-treated cockroaches compared to controls (Fig. 4A). Preliminary dissections of Myo-depleted cockroaches in N5D4 and N6D4 revealed PG size was greater than in controls (Fig. 4B). Measurement of the PG arm width during N5 and N6 showed that the PG from Myo-depleted cockroaches grew bigger than in the controls (Fig. 4C). Moreover, microscope examination of the PG at high magnifications using Edu labeling revealed that Myo-depleted cockroaches exhibited much more active cell division than controls (Fig. 4D).

**Fig. 4.**
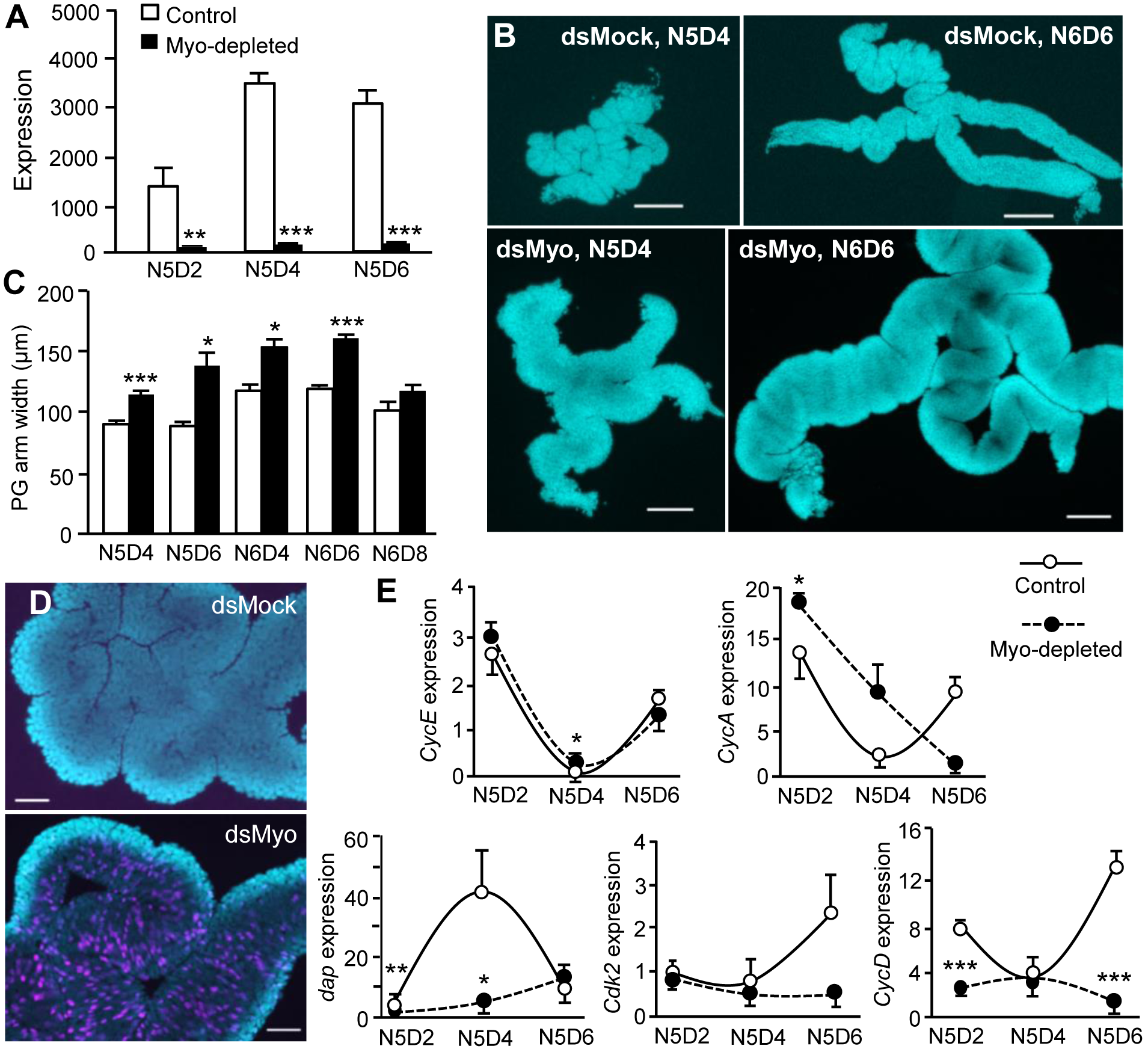
Effects of *myoglianin* mRNA depletion on prothoracic gland growth in *Blattella germanica*. (A) mRNA levels of *myo* in PG of Myo-depleted cockroaches and in controls, measured on 2-, 4- and 6-day old fifth instar female nymphs (N5D2, N5D4 and N5D6). (B) DAPI stained PGs from control (dsMock) and Myo-depleted (dsMyo) specimens in N5D4 and N6D6; scale bar: 200 μm. (C) PG arm width along different days of N5 (N5D4, N5D6), and N6 (N6D4, N6D6 and N6D8), comparing Myo-depleted cockroaches and controls. (D) Double labeling EdU (discrete pink spots) and DAPI (gland background bluish color) of PGs of N5D4, comparing Myo-depleted cockroaches and controls; scale bar: 50 μm. (E) mRNA levels of *CycE*, *CycA*, *dap*, *Cdk2* and *CycD* in PG tissues from dsMyo-treated cockroaches and controls on N5D2, N5D4 and N5D6. In A and E, results are indicated as copies of the given transcript per 1000 copies of BgActin-5c mRNA. In all quantitative diagrams, results are expressed as the mean ± SEM (n=3-5) and asterisks indicate statistically significant differences with respect to controls (*p<0.05, **p<0.01, ***p<0.001) according to the student’s *t*-test.

The transition from G1 into DNA replication (S phase) is crucial in determining whether to enter a new cell cycle, and four factors, Cyclin E (CycE), Cyclin A (CycA), Dacapo (Dap) and Cyclin-dependent kinase 2 (Cdk2), are key players in this transition. The accumulation of both CycE and CycA, which form complexes with Cdk2, promotes the transition to the S phase, whereas Dap negatively regulates this transition, mainly by sequestering the CycE/Cdk2 complexes (Barr et al., 2016; Bertoli et al., 2013). Moreover, in mouse C2C12 cells, myostatin (the Myo homolog in mammals, Lo and Frasch 1999) induces Cyclin D (CycD) degradation, thus resulting in cell cycle arrest (Yang et al., 2006). Based on these past findings, we measured the expression of *CycE*, *CycA*, *dap*, *Cdk2* and *CycD* in the PG from Myo-depleted and control cockroaches.

In controls, results indicated low levels of *dap* expression at the beginning of N5, high values in the middle, and low levels again at the end of the instar. Contrastingly, *CdK2*, *CycE*, *CycA*, and *CycD* expression showed an inverse pattern (Fig. 4E), which approximately corresponds to the dynamics of cell proliferation (Fig. 3).). *dap* expression in Myo-depleted cockroaches was generally quite dramatically reduced, whereas *CycA*, *CycD*, and *Cdk2* expression progressively decreased during N5. *Cyc*E expression was not significantly affected (Fig. 4E). Dap is a member of the mammalian p21 family of cyclin-dependent kinase inhibitors (CKIs) and inhibits the G1-to-S phase transition by sequestering the CycE/Cdk2 complex in a stable but inactive form (Lane et al., 1996). Induction of *dap* transcription causes a rapid accumulation of Dap protein, which inhibits CycE/Cdk2 activity and leads to G1 cell cycle arrest (Swanson et al., 2015). Conversely, *dap* knockdown leads to tissue hypertrophy and cancer processes (Kiyokawa et al., 1996; Nakayama et al., 1996). We propose that downregulation of *dap* expression may have maintained cell cycle activity in the PG of Myo-depleted cockroaches, thus leading to cell hyperproliferation and PG hypertrophy. The *CycE*, *CycA*, and *CdK2* expression patterns in Myo-depleted cockroaches could be a product of *dap* downregulation and the complex epistatic relationships between these factors (Barr et al., 2016; Bertoli et al., 2013). General downregulation of *CycD* expression in the PG of Myo-depleted cockroaches suggests that Myo has either a direct or indirect stimulatory effect on the expression of this cyclin. This effect differs from phenomena observed in mammals, where myostatin induces CycD degradation (Yang et al., 2006).

### Myoglianin depletion reduces the expression of ecdysone synthesis genes in the prothoracic gland

Myo depletion increased the duration of the fifth nymphal instar (Fig. 2A). To study the reasons for this delay, we therefore measured the expression of the following four genes, which code for ecdysone biosynthesis enzymes: *neverland* (*nvd*), which converts cholesterol to 7-dehydrocholesterol; *phantom* (*phm*) and *disembodied* (*dib*), which catalyze the addition of a hydroxyl group to the 25 and 22 carbon of the cholesterol side chain, respectively; and *shadow* (*sad*), which catalyzes the addition of a hydroxyl group to the 2 carbon of the cholesterol ring (Niwa and Niwa, 2014). Results showed that the expression of *nvd* was dramatically downregulated in N5D4 Myo-depleted cockroaches. The expression of *phm* and *dib* tended to be downregulated, whereas that of *sad* was unaffected (Fig. 5A).

**Fig. 5.**
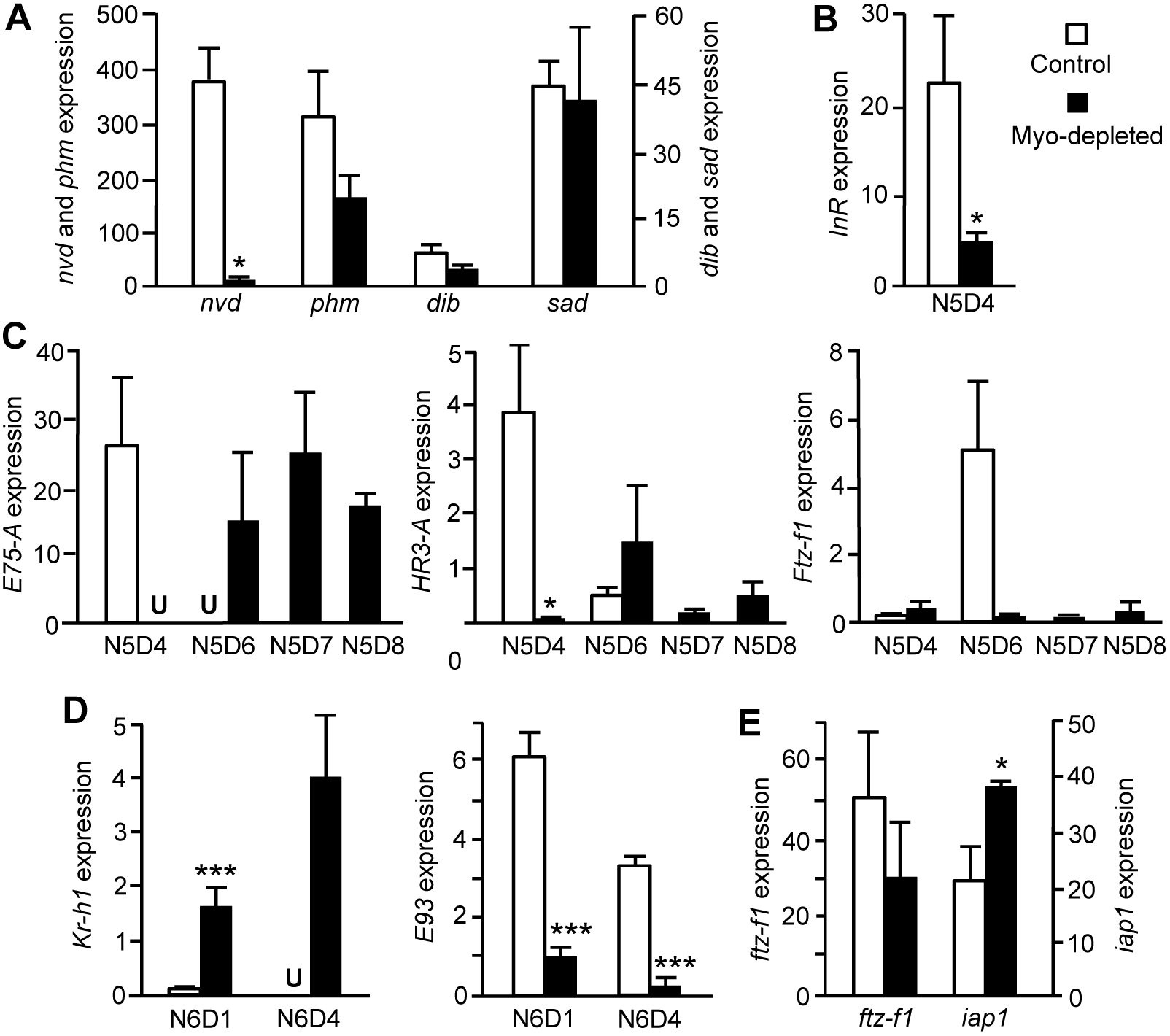
Effects of *myoglianin* mRNA depletion on the expression of prothoracic gland factors in *Blattella germanica*. (A) mRNA levels of *nvd*, *phm*, *dib* and *sad* in PG from Myo-depleted cockroaches and controls on 4-day-old fifth instar female nymphs (N5D4). (B) mRNA levels of *InR* in PG from Myo-depleted cockroaches and controls on N5D4. (C) mRNA levels of *E75-A*, *HR3-A* and ftz-f1 in PG from Myo-depleted cockroaches and controls, measured on different days of N5 (N5D4, N5D6 in controls, and N5D4, N5D6, N5D7 and N5D8 in dsMyo-treated). (D) mRNA levels of *Kr-h1* and *E93* in PG from Myo-depleted cockroaches and controls on N6D1 and N6D4. (E) mRNA levels of *iap1* and *ftz-f1* in PG from Myo-depleted cockroaches and controls on N6D8. Results, indicated as copies of the given transcript per 1000 copies of BgActin-5c mRNA, are expressed as the mean ± SEM (n=3-5); “U” in C and D diagrams means that the expression was under the limits of detection; asterisks indicate statistically significant differences with respect to controls (*p<0.05, ***p<0.001) according to the student’s *t*-test.

In *D. melanogaster*, the Activin branch of the TGF-β pathway promotes the expression of *nvd*, *dib* and *spookier* (*spok*, which is involved in catalyzing the conversion of 7-dehydrocholesterol to delta4, diketol), while it does not apparently affect *phm* and *sad* expression (Gibbens et al., 2011). These genes may be directly affected by the TGF-β pathway or this pathway’s influence could be mediated by the PTTH/torso pathway, acting at a transcriptional level, and the insulin pathway, acting post-transcriptionally (Gibbens et al., 2011). To date, there is no evidence that a PTTH/Torso pathway exists in cockroaches. Therefore, we focused on the Insulin pathway by measuring *Insulin receptor* (*InR*) expression, which was downregulated in the PG from Myo-depleted cockroaches (Fig. 5B). Incidentally, if Myo has a systemic stimulatory effect on *InR* expression, it could also explain the small size of the N6 instar emerging from dsMyo-treated N5 cockroaches (Figs. 2C and D).

We propose that Myo promotes *nvd* expression and, to a lesser extent, that of *phm* and *dib*. The stimulatory action on *nvd* and *dib* could be direct or mediated by the Insulin pathway, as in *D. melanogaster*. In the case of *B. germanica*, however, the Insulin pathway would act at a transcriptional level on these genes, rather than post-transcriptionally, as occurs in *D. melanogaster* (Gibbens et al., 2011). Additionally, we cannot rule out the possibility that the apparent stimulation of *phm* and *dib* transcription might also be mediated by FTZ-F1. Whereas FTZ-F1enhances *phm* and *dib* expression in the ring gland from late third instar larva of *D. melanogaster* (Parvy et al., 2005), our measurements revealed that *ftz-f1* expression was delayed and downregulated in the PG of Myo-depleted cockroaches (see below, Fig. 5C). This would support the hypothesis that FTZ-F1 helps enhance *phm* and *dib* transcription in the PG of *B. germanica*.

We also examined the expression of the ecdysone-dependent factors E75-A, HR3-A, and FTZ-F1, which have previously been characterized in *B. germanica* (Cruz et al., 2007; Cruz et al., 2008; Mané-Padrós et al., 2008). We noted that the expression pattern of these three genes in the PG of Myo-depleted cockroaches was conspicuously reduced (especially that of *ftz-f1*) and delayed with respect to controls (Fig. 5C). This effect is reminiscent of the developmental delay observed in *D. melanogaster* with impaired signaling in the Activin branch of the TGF-β pathway, which was ultimately due to deficient ecdysone production by the PG (Brummel et al., 1999; Gibbens et al., 2011). The same mechanism may explain why N5 lasts longer in Myo-depleted cockroaches (Fig. 2B).

### Myoglianin depletion prevents the onset of prothoracic gland degeneration

In the PG of *B. germanica*, ecdysone signaling promotes the degeneration of the gland towards the end of N6 and the beginning of adult life. This is mediated by FTZ-F1, a distal transducer in the ecdysone signaling pathway that is expressed at markedly high levels in the PG in the last day of N6 (N6D8) (Mané-Padrós et al., 2010). On the other hand, inhibitor of apoptosis-1 (IAP1) protects the PG from degeneration in early N6 and earlier nymphal stages. *iap1* shows maximal expression at N6D7 before it starts to decrease at N6D8, coinciding with a sharp peak in *ftz-f1* expression (Mané-Padrós et al., 2010). JH also plays a protective role, as treating N6 with the JH analog methoprene attenuates the expression of *ftz-f1* in N6D8 and prevents PG degeneration (Mané-Padrós et al., 2010). E93 may also be involved in PG degeneration in *B. germanica* as it is known to have pro-apoptotic functions, for example, in the degradation of salivary glands during *D. melanogaster* metamorphosis (Baehrecke and Thummel, 1995; Lee and Baehrecke, 2001; Lee et al., 2000).

In our experiments, Myo-depleted N6 cockroaches molted to a supernumerary N7, and, in some cases, successively on to N8 and adult, evidencing the PG was functional after N6. To study the mechanisms that prevented PG degeneration, we measured the expression of *Kr-h1*, as a key transducer of the JH signal, and *ftz-f1*, *iap1*, and *E93*, as potential regulators of the onset of PG degeneration. Results showed that *Kr-h1* was upregulated whereas *E93* was downregulated in N6 Myo-depleted cockroaches (Fig. 5D). This presumably results from the higher production of JH in the CA, where Myo-depleted cockroaches maintained upregulated *jhamt* expression in N6, with the consequent increase in *Kr-h1* transcription (Fig. 2E). Given that Kr-h1 represses *E93* expression in metamorphic tissues (Belles and Santos, 2014), *E93* was consequently downregulated in the PG (Fig. 5D). On the last day of N6 (N6D8), *ftz-f1* and *iap1*, respectively, showed the highest and lowest expression values within the instar (Mané-Padrós et al., 2010). In the PG of Myo-depleted cockroaches, however, *ftz-f1* expression in N6D8 tended to be lower than in controls (40% on average) whereas *iap1* expression was significantly upregulated (73% on average) (Fig. 5E). This is consistent with the notion that IAP1 and JH protect PG from degeneration, whereas FTZ-F1 plays a determinant pro-apoptotic role (Mané-Padrós et al., 2010).

## DISCUSSION

Transcriptome data indicated that *myo* expression is highly upregulated in N5 of *B. germanica*. This general upregulation in N5 is consistent with high expression levels measured by means of qRT-PCR at the beginning of the instar in the CC–CA and towards the mid-end of the instar in the PG.

Comparison of the expression levels for the measured factors suggests that high Myo levels in N5 repress *jhamt* expression in the CC–CA, so there is a reduction in JH production at the beginning of N6 (Fig. 6). In juvenile instars, JH acting through Kr-h1 represses *ftz-f1*expression and, in turn, FTZ-F1 represses *iap1* expression in the PG. In N6, low levels of circulating JH results in lower *Kr-h1* expression in the PG, with the concomitant de-repression of *ftz-f1* expression, inhibition of *iap1*, and activation of the caspase-mediated gland degeneration mechanisms (Fig. 6). As *E93*, a distal gene activated by ecdysone signaling (possibly by FTZ-F1), is an apoptosis triggering factor which is repressed by Kr-h1, it is plausible that the pro-apoptotic action of FTZ-F1 is mediated by E93 (Fig. 6).

**Fig. 6.**
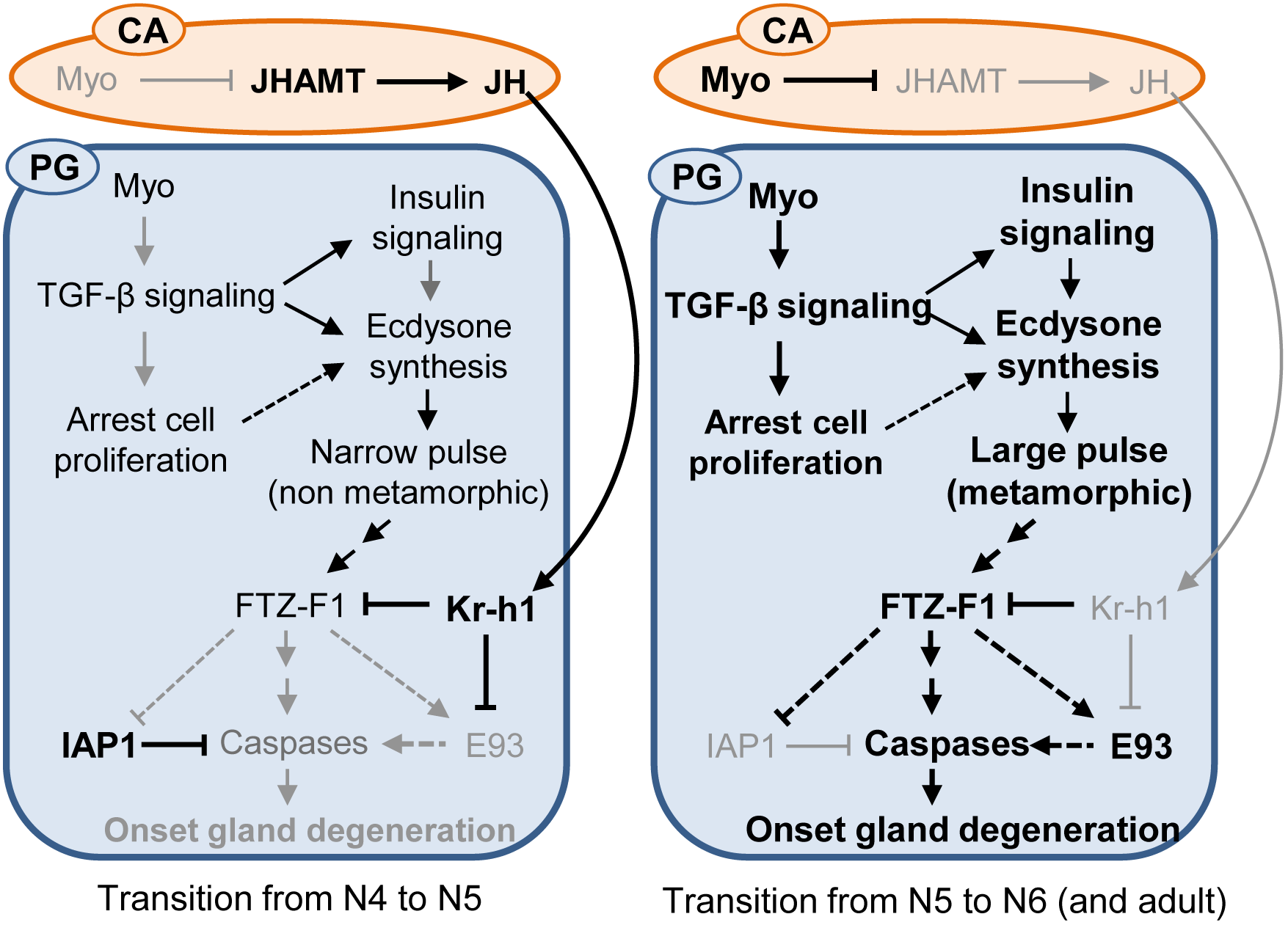
Roles of Myoglianin in the corpora allata (CA) and the prothoracic gland (PG) in the transition to the pre-metamorphic stage of *Blattella germanica*. The scheme of Myo roles in the transition from antepenultimate (N4) to penultimate (N5) nymphal instar is shown on the left, and the transition from N5 to the final nymphal instar (N6) and the adult on the right. High levels of the indicated factors and corresponding interactions and final processes, are indicated in bold, whereas low levels are indicated in grey. Continuous lines indicate interactions suggested by the experiments, whereas dashed lines indicate interactions hypothesized. See the text for details.

In the PG, *myo* is expressed during N5 and N6, but there is a marked peak of expression towards the mid-end of N5. Our data indicate that Myo enhances the expression of ecdysone biosynthesis genes, especially *nvd*, thus contributing to the production of the large, metamorphic ecdysone pulse in N6. The data also suggest that the peak of *myo* expression in N5 helps repress cell proliferation (at least in part by enhancing the expression of *dap*). Control over cell proliferation in N6 may be required to produce the subsequent ecdysone pulse (Fig. 6).

Our overall findings point to Myo as a key trigger of the transition from penultimate to last (pre-metamorphic) nymphal instar in *B. germanica*. The repressor role of Myo on *jhamt* expression and JH production has previously been reported in the cricket *G. bimaculatus* (Ishimaru et al., 2016). Although the functions of Myo in the PG were not examined in this insect, Ishimaru et al. (2016) noticed that *myo* expression levels were high in this gland. We presume, therefore, that the trigger role Myo plays in the pre-metamorphic stage of *B. germanica* may be extended to *G. bimaculatus*, and perhaps to other hemimetabolan insects. Intriguingly, the *myo* expression in *D. melanogaster* does not exhibit an especially prominent peak during the larval and pupa stages, neither in the Northern blot analyses published by Lo and Frasch (1999, figure 2O), nor in the transcriptome profiles reported by Ylla et al. (2018, figure 5E). Importantly, the larva–pupa–adult transition in the holometabolous metamorphosis requires a relatively rapid pattern of decreasing–increasing–decreasing JH production (Belles, 2011; Jindra et al., 2013). In this context, a dramatic interruption of JH production mediated by the repressive action of Myo on CA *jhamt* expression might not be suited for regulating metamorphosis in holometabolan species. Hence, we conclude that Myo’s role as a pre-metamorphosis trigger, at least in relation to the action in the CA and to JH, might be restricted to hemimetabolan insects, as this mechanism would has been lost in the evolutionary transition from hemimetaboly to holometaboly.

## MATERIALS AND METHODS

### Insects

The *B. germanica* cockroaches used in the experiments described herein were obtained from a colony fed on Panlab dog chow and water *ad libitum,* and reared in the dark at 29 ± 1 ^□^C and 60-70% relative humidity. Prior to injection treatments, dissections and tissue sampling, the cockroaches were anesthetized with carbon dioxide.

### RNA extraction and retrotranscription to cDNA

RNA extractions were carried out with the Gen Elute Mammalian Total RNA kit (Sigma-Aldrich, Madrid, Spain). A sample of 200 ng from each RNA extraction was used for mRNA precursors in the case of fat body, epidermis and muscle. All the volume extracted of CC–CA complex, prothoracic gland, brain, and ovary was lyophilized in the freeze-dryer FISHER-ALPHA 1-2 LDplus and then resuspended in 8 μl of milliQ H_2_O. RNA quantity and quality were estimated by spectrophotometric absorption at 260 nm in a Nanodrop Spectrophotometer ND-1000^®^ (NanoDrop Technologies, Wilmington, DE, USA). The RNA samples then were treated with DNase (Promega, Madison,WI, USA) and reverse transcribed with first Strand cDNA Synthesis Kit (Roche) and random hexamers primers (Roche).

### Determination of mRNA levels by quantitative real-time PCR

Quantitative real-time PCR (qRT-PCR) was carried out in an iQ5 Real-Time PCR Detection System (Bio-Lab Laboratories), using SYBR^®^Green (iTaq^TM^ Universal SYBR^®^ Green Supermix; Applied Biosystems). Reactions were triplicate, and a template-free control was included in all batches. Primers used to detect mRNA levels studied here are detailed in Table S1. We have validated the efficiency of each set of primers by constructing a standard curve through three serial dilutions. In all cases, levels of mRNA were calculated relative to BgActin-5c mRNA (accession number AJ862721). Results are given as copies of mRNA of interest per 1000 or per 100 copies of BgActin-5c mRNA.

### RNA interference

The detailed procedures for RNAi assays have been described previously (Ciudad et al., 2006). Primers used to prepare BgMyo dsRNA are described in Table S1. The sequence was amplified by PCR and then cloned into a pST-Blue-1 vector. A 307 bp sequence from *Autographa californica* nucleopolyhedrosis virus (Accession number K01149.1) was used as control dsRNA (dsMock). A volume of 1 μl of the dsRNA solution (3μg/μl) was injected into the abdomen of freshly emerged fifth instar female nymphs (N5D0), with a 5μl Hamilton microsyringe. Control insects were treated at the same age with the same dose and volume of dsMock.

### Morphological studies of prothoracic glands

The PGs were studied in female nymphs at chosen ages and stages. The glands were dissected out of the first thoracic segment of the animal under Ringer’s saline. The glands were fixed in 4% paraformaldehyde in PBS for 1h, then washed with PBS 0.3% Triton (PBT) and incubated for 10 min in 1 mg/ml DAPI in PBT. They were mounted in Mowiol (Calbiochem, Madison, WL, USA) and observed with the fluorescence microscope Carl Zeiss – AXIO IMAGER.Z1.

### EdU experiments to measure cell proliferation in prothoracic glands

EdU (5-ethynyl-2′-deoxyuridine) is a thymidine analogue recently developed for labeling DNA synthesis and dividing cells in vitro (Chehrehasa et al., 2009), which is more sensitive and practical than the commonly used 5-bromo-2′-deoxyuridine, BrdU. We followed an approach in vivo, using the commercial EdU compound “Click-it EdU-Alexa Fluor^®^ 594 azide” (Invitrogen, Molecular Probes), which was injected into the abdomen of the nymphs at chosen ages and stages with a 5μl Hamilton microsyringe (1 μl of 20 mM EdU solution in DMSO). The control specimens received 1 μl of DMSO. PGs from treated specimens were dissected 1 h later, and processed for EdU visualization according to the manufacturer’s protocol.

## Acknowledgements

We thank Guillem Ylla and Jose Carlos Montañes for helping with transcriptome comparisons and searching for genes in the *B. germanica* genome, available at https://www.hgsc.bcm.edu/arthropods/german-cockroach-genomeproject, as provided by the Baylor College of Medicine Human Genome Sequencing Center. We also thank Alba Ventos-Alfonso for helping in different experiments and image treatment, and Maria-Dolors Piulachs and Jose-Luis Maestro for helpful discussions.

## Author Contributions

X.B. conceived the study. O.K. performed the RNAi experiments and the qRT-PCR measurements. X.B. and O.K. discussed and interpreted the results, and commented on the manuscript. O.K. drafted a first version of the manuscript. X.B. wrote the final version of the manuscript.

## Data availability

Whole-transcriptome profiling data are available at GEO with the Accession Number GSE89295.

## Supplementary information

Supplementary information available online at

